# Synlet: an R package for systemically analyzing synthetic lethal RNA interference screen data

**DOI:** 10.1101/043570

**Authors:** Chunxuan Shao, Frank Westermann, Thomas Höfer

## Abstract

**Summary:** High-throughput synthetic lethal RNA interference (RNAi) screen experiments shed important insights on the filed of cancer researches and drug discovery, but a comprehensive software for analyzing the data was not available yet. We present synlet, an R package provided a complete pipeline to process the synthetic lethal RNAi screens data. Synlet provides several methods to access the screen quality, including Z’ factor and data visualization. B-score and fraction of control or samples **normalization** methods are implemented in the package. More importantly, synlet facilitates the process of hits selection by implementing several algorithms, providing the possibility to identify high confidence targets.

**Availability:** The source code is freely available in Bioconductor (http://bioconductor.org/).

## 1 Introduction

The RNA interference (RNAi) technique allows knockdown of specific genes and quickly becomes a vital tool in exploring biological questions. Recent developments in RNAi methodology combine silencing of a single gene together with the genomeic methods to make it feasible to target the complete genome (Mohr *et al*., 2014). For example, high-throughput RNAi screens could be performed in 384-wells microplates, in which each well may contain reagents (for example, small interfering RNA (siRNA)) designed for a specific gene. An interesting extension of the RNAi screen is the synthetic lethal screens that focus on identifying genes lead to higher mortality rate under certain conditions or genetic mutations, shedding important insights on the fields of cancer research as it provides an straightforward approach to identify genes mainly support the growth of cancer cells (Ngo *et al*., 2006; Schlabach *et al*., 2008; Luo *et al*., 2009).

Many tools are available in handling the high-throughput RNAi screens data. CellHTS (Boutros *et al*., 2006) and RNAither (Rieber *et al*., 2009) are R packages that provide comprehensive pipelines for the screen data from quality accessment to hits selection. ScreenSifter (Kumar *et al*., 2013) provides a user-friendly interface to manage multiple

RNAi screen projects. HTSanalyzeR (Wang *et al*., 2011) focuses on enrichment and network analysis of screen results by combining several well known software. However, to the best of our knowledge, there is no software provided an automated pipeline for analyzing the synthetic lethal RNAi screen data.

Here we present synlet, an R package building a complete analysis pipeline for synthetic lethal RNAi screen data. Synlet provides various ways to check RNAi screen quality, including the widely used Z’ factor (Zhang, 1999) and several visualization tools. The raw screen data could be normalized by the B-scores method (Malo *et al*., 2006), fraction of control or fraction of samples (Birmingham *et al*., 2009). Finally, synlet allows to select genes with synthetic lethal effect by several algorithms, including Student’s t-test, median ± k median absolute deviation (Chung *et al*., 2008), rank products (Breitling *et al*., 2004; Hong *et al*., 2006) and redundant siRNA activity (König *et al*., 2007). A scheme described the analyzing pipeline is shown in Figure 1, which includes four major steps (S1 to S4) and major functions implemented.

**Fig. 1.**
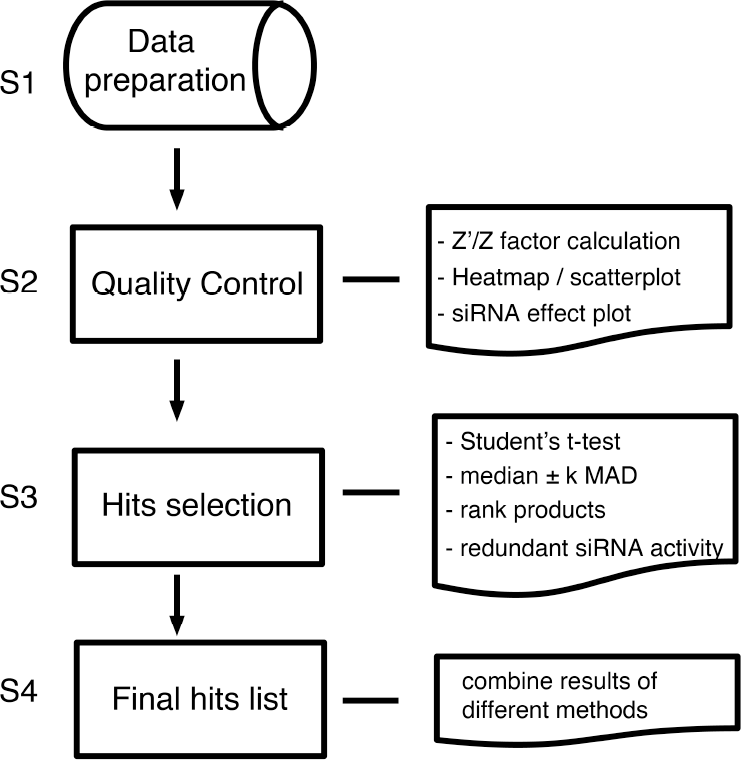
Synthetic RNAi analysis pipeline. The scheme describes the working flow and major functions in synlet.

## 2 Package description

### 2.1 Input Data format

Synlet requires the screen data stored in the format of data frame, which is simple to generate and manipulate. Eight columns are mandatory for the data, including “PLATE”, “MASTER_PLATE”, “WELL_CONTENT_NAME”, “EXPERIMENT_TYPE”, “EXPERIMENT_MODIFICATION”, “ROW_NAME”, “COL_NAME” and “READOUT”. “PLATE” is the basic screen unit in the experiments, several control and treatment “PLATE”s construct a “MASTER_PLATE”. The hits selection steps are usually performed based on “MASTER_PLATE”. “WELL_CONTENT_NAME” shows the siRNA id and targeted genes in each well. The microplate wells containing negative or positive control siRNA are specified in “EXPERIMENT_TYPE”. The treatment and control plates are labeled in “EXPERIMENT_MODIFICATION”. An example dataset is included in the R package.

### 2.2 Quality control and data visualization

There are several ways to check the RNAi screen quality in synlet. Z factor and Z’ factor (Zhang, 1999), which are commonly used in quality accessment, could be easily calculated for each plate. In addtion, the data visualization play a important role in identifying possible technical variations, like the edge effect or failures of negative or positive control siRNAs. Synlet implemented several functions to plot RNAi screen data. It is feasible to plot the raw RNAi screen data of all plates as a heatmap with the same layout as the experiments, providing an overview of data quality (Supplementary Figure S1). The scatter plots are employed to represent the screen output of all plates by colorful dots, highlighting the effect of control siRNAs (Supplementary Figure S2). Further, synlet could plot the raw and normalized screen results of a single gene in the control and treatment plates together with Z’ factor, which is helpful to examine the knockdown effect of interested genes in detail (Supplementary Figure S3).

### 2.3 Normalization of data

The purpose of normalization is to remove unwanted technical variations in screen experiments and prepare the data for the downstream analysis. Proper normalization of raw output is an important step in correctly interpreting the screen results. Synlet provides different methods to normalize the data: the B-score approach, fraction of control and fraction of samples. B-scores are robust normalized value against row and column bias generated by the two-way median polish algorithm and division of adjusted median absolute deviation (Malo *et al*., 2006). In fraction of control and fraction of samples normalization, raw output of wells are divided by the mean or median of control siRNAs and whole plate output, respectively (Birmingham *et al*., 2009).

### 2.4 Hits selection

The main goal of sythentic lethal RNAi screen experiements is to identify genes (hits) led to reliable difference in mortality between treatment and control plates. It could be a difficult task because of cell heterogeneity, reagent efficiency, etc. Synlet tries to improve the results of hits selection by employing several algorithms that explore knockdown effect from different directions, including student’s t-test, median ± k median absolute deviation (Chung *et al*., 2008), rank products (Breitling *et al*., 2004) and redundant siRNA activity (RSA) (König *et al*., 2007).

Student’s t-test is commonly used to test whether the mean from two samples are identical, thus it could be a helpful strategy in identifying synthetic lethal genes (Whitehurst *et al*., 2007). The B-scores calculated from control and treatment plates are used since they are robust to outliers in the data. The multiple comparison problem is addressed by the Benjamini and Hochberg method (Benjamini and Hochberg, 1995).

Hits selection based on median ± *k* median absolute deviation (MAD) is an recommended appraoch in RNAi screen data analysis due to the easy calculation and robustness to outliers revealed in the real data and simulation study (Chung *et al*., 2008). Normalized screen outputs that are *k* MADs away from median of the plate provide evidences for hits selection. Synlet calculates the median and MAD based on the ratio between treatment and control plates, where raw data are normalized by fraction of control or fraction of samples normalization. By default *k* is set to be three.

The rank products algorithm is a non-parametric statistic method proposed to find consistently up/down-regulated genes between treatment and controls replicates in microarray data and has been successfully used in analyzing the RNAi screen data (Rieber *et al*., 2009). It has several advantages over the parametric Student’s t-test, including clear biological meaning, fewer assumptions of the data and improved performance. Synlet uses the rank products method to compare the normalized screen results between treatment and control plates and identified siRNAs accelerating or hampering cell growth. *P*-value, false positive predictions and fold changes are provided to facilitate hits selection.

Redundant siRNA activity (RSA) is a distiguished method proposed for RNAi screen data that systemically employs the information provided by multiple siRNAs targeting a single gene to reduce the off-target and improve confirmation rate. Briefly, RSA calculates a *P*-value for the rank distribution of multiple siRNAs silenced the same gene under the background of all siRNA signals in the experiment by iterative hypergeometric distribution formula (König *et al*., 2007). Compared to the methods mentioned above, siRNAs targeted the same genes have identical P-value, and genes with several moderately effect siRNAs may have smaller P-value than genes with fewer strong effect siRNAs. Synlet provides a wrapper function to use the RSA R codes and applies the algorithm to the ratio between treatment and control signals normalized by fraction of control or fraction of samples.

## 3 Conclusion

The synlet package provides an all-in-one toolbox to analyze synthetic lethal RNAi screen data from quality control and visualization to hits selection. Several tools are implemented in the package to display the complete experiment signals and individual gene single. Synlet emphasizes the hits selection step, in which several methods with different statistic property are included. This package will likely facilitate the analysis of RNAi screen data and shed insights on cancer researches and drug discovery.

## Acknowledgements

We thank Sina Gogolin for helpful discussions, Dr.Yingyao Zhou for kindly providing the RSA codes.

